# Vitamin D3 Deficiency Exacerbates Abdominal Aortic Aneurysm Progression Via Complement C3a Activation

**DOI:** 10.64898/2026.06.05.730431

**Authors:** Aravinthan Adithan, Jospeh B. Hartman, Walker Ueland, Jeff Arni C. Valisno, Gang Su, Michael Fassler, Shiven Sharma, Carl Atkinson, Jennifer K. Mulligan, Ashish K. Sharma, Gilbert R. Upchurch

## Abstract

Abdominal aortic aneurysm (AAA) is a chronic inflammatory vascular disease characterized by progressive extracellular matrix degradation, vascular smooth muscle cell (VSMC) loss, and immune cell infiltration, ultimately leading to aortic dilation and rupture. Although vitamin 25(OH)D_3_ deficiency has been associated with cardiovascular inflammation, it’s mechanistic role in AAA pathogenesis remains poorly defined. Here, we investigated the role of vitamin D□ mediated signaling to regulate complement pathway activation, particularly the C3a axis, to modulate aneurysm development. Single cell-RNA sequencing analysis of human tissue demonstrated significant differences in Vitamin D and complement pathway-related genes in VSMCs in AAAs compared to control aortic tissue. Using a murine elastase-induced AAA model, we observed that vitamin D_3_–deficient diet significantly enhances aortic dilation, leukocyte infiltration, proinflammatory cytokine expression and elastin fragmentation, as well as decreases SMC α-actin expression compared with vitamin D3–sufficient conditions. Furthermore, vitamin D3 deficiency was accompanied by increased aortic expression of complement component C3a that correlated with vascular inflammation and remodeling during AAA progression. Pharmacological blockade with a C3a receptor antagonist (C3aRA) markedly attenuated AAA formation in two established murine AAA models with concomitant reductions in proinflammatory cytokines and preservation of aortic wall structure. *In vitro* studies demonstrated that stimulation of VSMCs significantly increased C3a production, which was suppressed by calcitriol (active form of Vitamin D) treatment. These studies suggest that the vitamin D–C3a axis is a critical regulator of vascular inflammation and AAA progression, and postulate that restoring vitamin D□ sufficiency or targeting C3a signaling may represent a novel therapeutic strategy to limit AAA growth and rupture.

**Graphical Abstract:** 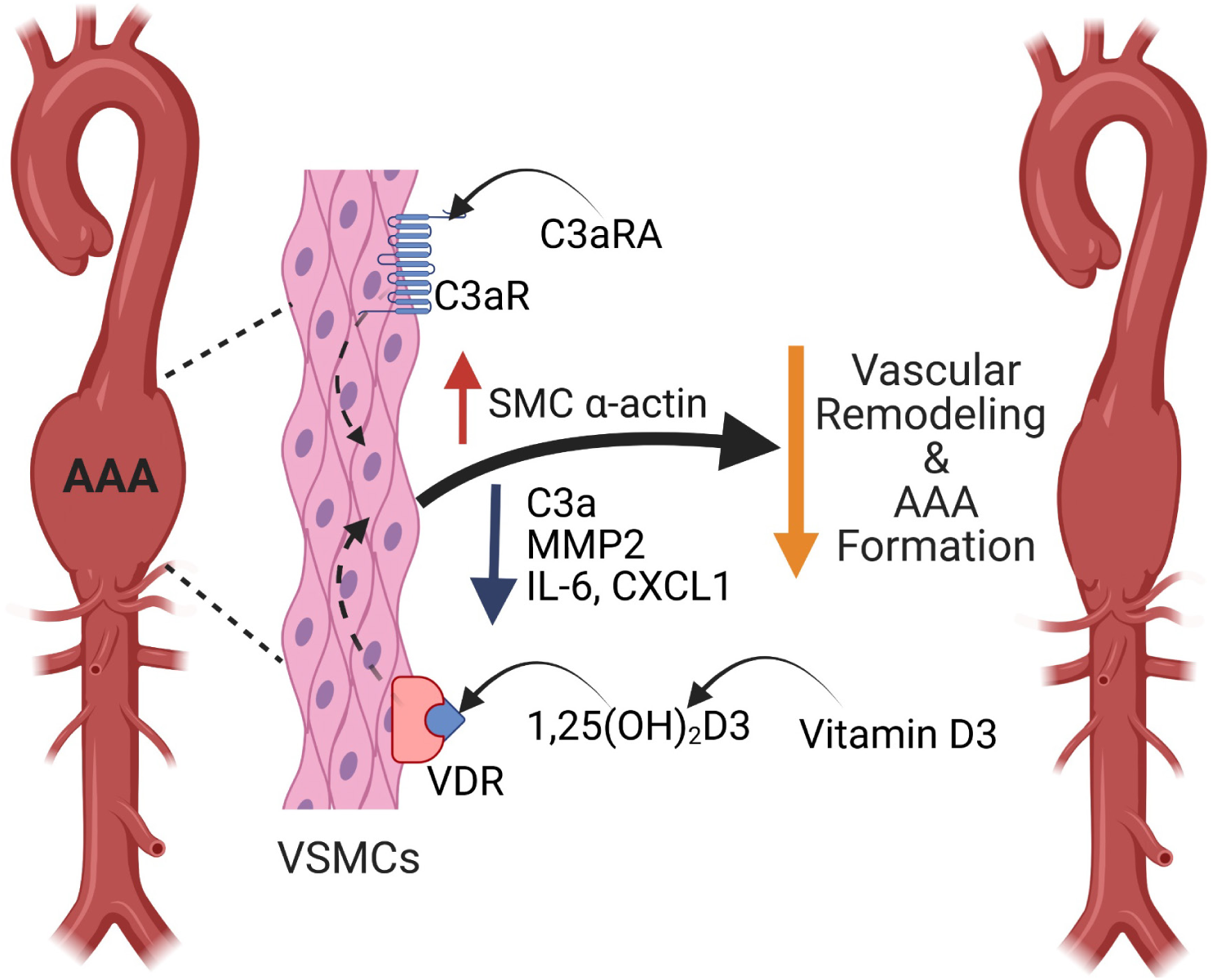

## INTRODUCTION

Abdominal aortic aneurysms (AAA) are a life threatening vascular disorder characterized by progressive weakening and dilation of the aortic wall, associated with extracellular matrix (ECM) degradation and vascular smooth muscle cell (VSMC) dysfunction. This persistent inflammatory milieu can lead to rupture resulting in up to an 81% mortality rate among men (1). Chronic inflammation within the aortic wall, characterized by the infiltration of neutrophils, macrophages, and lymphocytes, along with the activity of proteolytic enzymes, is regarded as a principal mechanism underlying aneurysm progression and medial degeneration (2). Recent studies have postulated that complement pathway activation is a key mediator of pathological vascular remodeling, contributing to lipid/protein deposition and modification within the vessel wall and inciting a robust immune inflammatory response (3). Moreover, complement serum levels have been reported to be elevated in patients with AAA (4).

VD_3_ is a secosteroid hormone, which has been shown to be synthesized in the skin, where pro-vitamin D_3_ is metabolized to pre-vitamin D_3_ followed by binding to vitamin D binding protein, and transported to the liver where it is metabolized to 25-hydroxycholecalciferol [25(OH)D_3_] (5, 6). The hormonally active isoform is then generated via 1α-hydroxylase that converts 25(OH)D_3_ to its active form, 1α,25-dihydroxyvitamin D_3_ [1,25(OH)_2_D_3_] which mediates its actions through the vitamin D receptor (VDR) expressed ubiquitously in vascular cells (7). The active hormonal form of vitamin D, 1-α,25-dihydroxyvitamin D3, extends its role beyond calcium homeostasis and bone metabolism to the regulation of immune responses, vascular cell function, and inflammation (7, 8). Both epidemiological and experimental studies implicate vitamin D deficiency and impaired vitamin D signaling in adverse cardiovascular outcomes, endothelial dysfunction, VSMC phenotypic switching, and enhanced inflammatory activation—processes central to aneurysm pathogenesis (9, 10). Although large cohort data on circulating vitamin D markers and incident AAA have been mixed, mechanistic work indicates that vitamin D can modulate cytokine expression, MMP activity and VSMC survival integral for modulating aortic wall integrity (8, 11). More importantly, low levels of 25(OH)D_3_ is associated with larger aortic aneurysm diameters and an increased risk of rupture, potentially due to elastin degradation via increased matrix metalloproteinase activity (12).

AAA progression depends on the interplay of inflammatory/proteolytic processes and anti-inflammatory signals wherein endogenous generation of active forms of vitamin D3 (1,25-dihydroxyvitamin D_3_ or calcitriol) plays a significant, though paradoxical, role in aortic aneurysms(13, 14). Conversely, activation of complement system in the aneurysm milieu dysregulates vitamin D signals, tilting the local environment toward unchecked inflammation and matrix breakdown. Early clinical and proteomic studies of AAAs have documented local complement retention with altered systemic complement levels and suggest complex spatial dynamics of complement proteins within intraluminal thrombus and aortic wall, supporting the biological plausibility of such localized interactions (15). Supporting the functional relevance of C3 in aneurysm biology, genetic and pharmacologic inhibition of C3-derived signaling reduces aortic inflammation and aneurysm formation in experimental models, indicating that interrupting C3 signaling can modify disease trajectory (16, 17). Together with data implicating dysregulated vitamin D activation/inactivation enzymes in immune cells and vascular cells, these observations motivate the hypothesis that complement C3 is a novel regulator of local vitamin D metabolism in the aneurysmal aortic wall acting as a regulatory axis that could amplify inflammation, impair efferocytosis, and accelerate matrix degradation leading to aneurysm growth and rupture.

In this study, we delineated the role of vitamin D_3_ deficiency in exacerbating AAA formation via complement C3 activation in vascular smooth muscle cells. Our data suggests that vitamin D and complement pathway-dependent genes are differentially regulated during human AAA formation. Furthermore, animals with Vitamin D_3_ deficient diet demonstrate exacerbated aortic diameter, aortic inflammation, C3a expression and vascular remodeling in murine AAAs whereas treatment with a C3a receptor antagonist (C3aRA)significantly attenuates aortic diameter, proinflammatory cytokine expression, matrix metalloproteinases (MMP2) expression and restores vascular integrity to decrease AAA formation and prevent aortic rupture. In summary, modulating the vitamin D_3_/C3a signaling pathway offers a novel and promising approach for attenuating AAA progression by enhancing inflammation-resolution and inhibiting SMC-dependent remodeling.

## MATERIALS AND METHODS

### Human Single Cell RNA Sequencing

Single cell RNA-seq data set of human AAAs and controls was obtained from Gene Expression Omnibus (GSE166676) and re-analyzed via Seurat(18). Cell annotations were assigned by extracting cluster specific markers via FindMarkers and cross-referencing the marker set with published human and mouse cell atlases via PanglaoDB(19). Cell annotations were verified via expression of putative cell markers for vascular smooth muscle cell (VSMC) clusters as described in previously published analyses of scRNA datasets of the aorta(20, 21). Differential gene expression in VSMC clusters between human AAA and controls were performed via the FindMarkers function in Seurat and statistical significance was assessed using the Wilcoxon Rank Sum Test. Differentially expressed genes (DEGs) were identified as genes with adjusted p-value < 0.05 and |fold change| >0.1 A list of vitamin D and complement-receptor related genes were generated using GeneCards (query=vitamin D or complement) and genes with relevance score >1 were cross-references with list of DEGs to identify differentially expressed vitamin D and complement-elated genes (DE-VDCRGs).

### Murine AAA Models

C57BL/6 (wild-type, WT) male mice, aged 8-12 weeks old obtained from Jackson Laboratory (Bar Harbor, ME, USA). Mice were maintained on a 12-h light cycle in a temperature-controlled room (25°C). All animal experimentation was approved by the University of Florida Institutional Animal Care and Use Committee (protocol # 201910902). Mice were anesthetized using isoflurane, and the infrarenal abdominal aorta was exposed through a midline laparotomy, as previously described (22). The aorta was carefully dissected circumferentially from surrounding tissues and topically treated for 5 minutes with 5 μL of elastase (0.4 U/mL, type I porcine pancreatic elastase (Sigma-Aldrich, St. Louis, MO) or heat-inactivated elastase (HIE), which served as a control. WT mice underwent AAA induction surgery and were treated with C3aRA (SB290157; Sigma Aldrich) administered at a concentration of 1 mg/kg/day via intraperitoneal injection. The C3aRA was administered from postoperative days 7 to 13, followed by aortic tissue harvest on day 14. In a separate cohort of the chronic AAA and aortic rupture model, C3aRA was administered from days 14 to 27, and aortas were harvested on day 28. Control mice group received equivalent volumes of sterile saline on matching schedules.

In the AAA and aortic rupture model, elastase application was combined with supplementation of 0.2% β-aminopropionitrile (BAPN) in drinking water continued through day 28 post-surgery, as previously described (23). Treated aortic sections were harvested post operatively on day 14 (topical elastase model) and day 28 (AAA and aortic rupture model), and preserved in paraformaldehyde for immune histochemistry or snap-frozen in liquid nitrogen for molecular analysis. Aortic diameter was measured using video micrometry (AmScope, Irvine, CA). The degree of aortic dilation was calculated using the formula: [(maximal aortic diameter − baseline diameter)/ baseline diameter] × 100. A dilation of ≥100% was considered indicative of AAA formation.

### Vitamin D3 diet regimen in murine elastase AAA model

WT mice were randomized into two dietary groups; Vitamin D3-deficient diet (Harlan Teklad TD.89123, Madison, WI) and Vitamin D3-sufficient control diet (TD.89124). Mice were maintained on their respective diets for 4 weeks prior to AAA induction surgery and continued the same diet until end of the experiments. Serum 25-hydroxyvitamin D3 [25(OH)D3] levels were measured using a commercial ELISA kit (ImmunoDiagnostic Systems, Fountain Hills, AZ) to confirm the vitamin D3 deficiency. At the endpoint, aortic tissues were collected, samples were stored at −80 °C for molecular analysis or fixed in 10% buffered formalin for histological evaluation.

### Histology

Harvested aortic tissues were fixed overnight in 10% zinc-buffered formalin, followed by dehydration through a graded ethanol series and embedding in paraffin. For immunohistochemical analysis, tissue sections were stained using the following primary antibodies: anti-mouse Mac-2 for macrophages (1:5000; Cedarlane Laboratories, Burlington, ON, Canada; catalog no. CL8942AP) and anti-mouse α-smooth muscle actin (α-SMA, 1:1000; Sigma, St. Louis, MO; catalog no. A5691). Verhoeff–Van Gieson (VVG) staining for elastin (Polysciences, Inc., Warrington, PA; catalog no. 25089-1) was quantified by counting the number of elastin fiber breaks per mm² using Fiji (ImageJ, NIH), and results were averaged and presented graphically. Imaging was performed at 20× magnification using a Nikon microscope equipped with a digital camera and NIS-Elements BR software. Quantification of staining intensity was performed using QuPath version 0.5 (QuPath; University of Edinburgh, Edinburgh, UK) by measuring the percentage of positively stained area relative to the total aortic cross-sectional area. Histological evaluation was performed on three separate aortic sections per animal, and quantification was conducted independently by blinded observer. Representative images from multiple animals per group were selected for consistency and accuracy

### mRNA expression analysis by quantitative RT-PCR

mRNA expression of C3, C3aR, C5 and C5aR in aortic tissue were evaluated by RT-PCR on day 14 in experimental groups using respective primer sequences (Supplementary Fig. S1). Total RNA was isolated from tissue samples using TRIzol reagent (Invitrogen) according to the manufacturer’s protocol. RNA purity and concentration were determined spectrophotometrically. One microgram of RNA was reverse transcribed using the iScript™ cDNA Synthesis Kit (Bio-Rad) in a 20 µL reaction volume. The resulting cDNA was diluted 1:10 with nuclease-free water for subsequent qPCR analysis. All primers were synthesized by Integrated DNA Technologies. Reactions were performed in a final volume of 20 µL containing 10 µL of 2× SYBR Green Master Mix, 0.4 µL of each primer (10 µM), 2 µL of cDNA template, and 7.2 µL of nuclease-free water. Thermal cycling conditions were as follows: 95 °C for 3 min, followed by 40 cycles of 95 °C for 15s and 60 °C for 30s. Relative mRNA expression was calculated using the comparative Ct 2^(−ΔΔCt) method, with GAPDH used as the endogenous control.

### Cytokine Multiplex Assay

Cytokine levels in murine aortic tissue homogenates were quantified using the Bio-Plex Bead Array system with a multiplex cytokine panel assay (Bio-Rad Laboratories, Hercules, CA), following the manufacturer’s instructions. Briefly, aortic tissues from various experimental groups were snap-frozen in liquid nitrogen, ground using a mortar and pestle, and homogenized in tissue extraction buffer. Protein concentration was determined using the BCA assay kit (Thermo Fisher Scientific), and equal amounts of protein were used for cytokine measurements in the tissue samples.

### *In vitro* experiments

Human primary vascular smooth muscle cells (HVSMCs; Lifeline, Cell Technology, SanDiego, CA) were cultured in VascuLife® basal medium supplemented with 10% charcoal-stripped fetal bovine serum (FBS; Gibco) and maintained at 37 °C in a humidified atmosphere containing 5% CO□. Upon reaching approximately 80% confluence, the culture medium was replaced with VascuLife® basal medium containing 1% charcoal-stripped FBS for 12–16 h prior to experimental treatments. Cells were pretreated with calcitriol (10 nM; Tocris Bioscience) for 24 h prior with or without porcine pancreatic elastase (0.4 U/mL; Sigma-Aldrich) for 5 min at 37 °C. Cells were then washed twice with 1x PBS and incubated with cytomix consisting of interleukin-17 (IL-17A), tumor necrosis factor-α (TNF-α), and high-mobility group box 1 (HMGB1), 50 ng/mL each, in basal medium supplemented with 1% charcoal-stripped FBS. After incubation, cell culture supernatants were collected, centrifuged at 1,500g for 10 min to remove cellular debris, and stored at −80 °C until further analysis. For RNA isolation, cells were lysed directly in the culture wells by the addition of 1 mL TRIzol™ reagent (Invitrogen) per well, and samples were stored at −80 °C until RNA extraction. Levels of human complement C3a were quantified in culture supernatants and in human aortic tissue samples (control and AAA) using a commercially available human C3a ELISA kit (Abcam, ab279352), according to the manufacturer’s instructions.

### Zymography assay

Matrix metalloproteinase 2 (MMP2) activity in aortic tissue and cell culture samples was assessed using the Novex™ Zymogram Plus Gelatin Gel Electrophoresis system (Thermo Fisher Scientific), following the manufacturer’s protocol. Briefly, equal amounts of protein extracted from aortic tissue homogenates were mixed with non-reducing sample buffer and loaded onto 10% gelatin-containing polyacrylamide gels. After electrophoresis, the gels were renatured and incubated in development buffer to allow enzymatic digestion of the gelatin substrate by active MMP. Gels were then stained with Coomassie Brilliant Blue and destained to visualize zones of proteolysis, which appeared as clear bands against a blue background. Both the pro- and active forms of MMP-2 were identified based on their molecular weights. Band intensity was quantified using ImageJ software.

### Statistical analysis

Statistical analysis was performed using GraphPad Prism 10 (GraphPad Software, La Jolla, CA) software. For comparisons involving three or more groups, a one-way ANOVA followed by Tukey’s multiple comparisons test was utilized. Pairwise comparisons were conducted using either an unpaired Student’s t-test or, a non-parametric Mann-Whitney test. Data are presented as mean ± standard error of the mean, and statistical significance was defined as p<0.05.

## RESULTS

### Single-cell RNA-sequencing analysis reveals dysregulation of Vitamin D/Complement signaling-related genes in AAA patients

To evaluate vitamin D and complement related pathway-mediated signaling in human AAAs, single-cell RNA sequencing data from aortic tissue from AAA patients and comparative controls was analyzed using a previously reported sequencing datasets GSE226492 and GSE152583 from Gene Expression Omnibus. Data was analyzed via Seurat and the VSMC cluster was isolated. Differential expression analysis comparing VSMCs from human AAAs (n=7) compared to control aortas (n=5) was performed via FindMarkers calculated using Wilcoxon Rank Sum Testing (20), and differentially expressed genes were identified (Fig. 1A). Data was analyzed via Seurat and the VSMC clusters were identified for further analyses (Fig 1B). A total of 11,641 genes were detected within the VSMC cluster and evaluated for differential expressions, wherein 716 genes were downregulated, 2297 genes were upregulated, and 8628 genes were unchanged (Supplementary Table S1). 16/716 downregulated genes and 57/2297 upregulated genes were related to vitamin D and complement related genes (VDCRGS) (Fig. 1C and 1D). Notably, this analysis suggests that genes known to regulate inflammation (TNF, IL6, NFkB, TGFB1, IL1B, APOE) were markedly dysregulated in AAAs (Fig. 1E). Moreover, the expression of complement C3a in aortic tissue of human AAA patients was significantly enhanced compared to controls (Supplementary Fig. S2). These findings in human AAA tissue indicate the association between vitamin D/complement-mediated signaling in aortic inflammation, vascular remodeling and cell death and supported further exploration of this pathway’s role in preclinical models of AAA.

**Figure 1.**
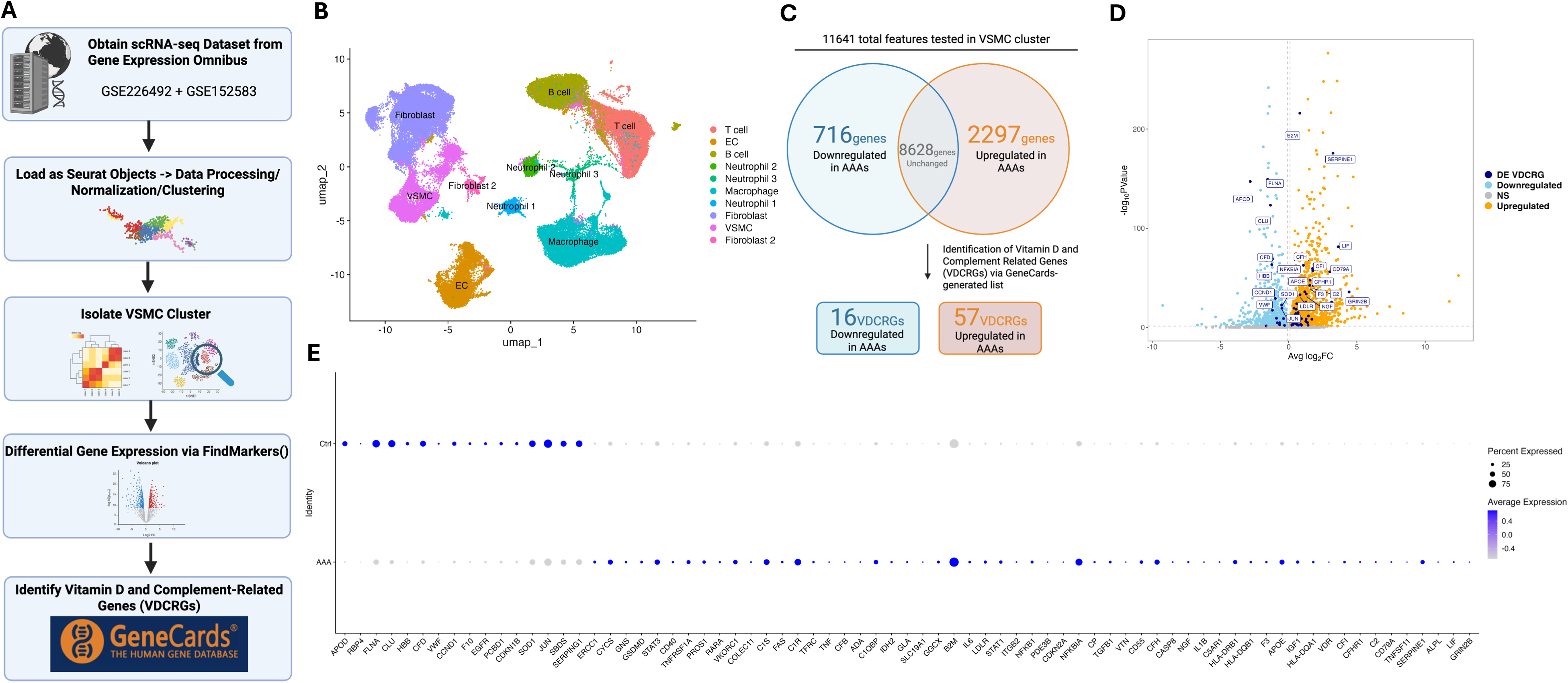
Vitamin D and complement-related genes are dysregulated in VSMCs of human AAAs. **A.** Bioinformatic workflow to analyze the human AAA scRNA-seq data sets. Differential expression analysis comparing VSMCs from human AAAs (n=7) vs control aortas (n=5) was performed via FindMarkers calculated using Wilcoxon Rank Sum Testing. Genes with adjusted p-value < 0.05 and |fold-change| > 0.1 were considered significant (DEGs). A list of Vitamin D and Complement related genes (VDCRGs) was generated using GeneCards and used to identify differentially expressed VDCRGs (DE VDCRGs). **B.** Uniform manifold projection (UMAP) plot of the annotated clusters from human AAAs and control. The VSMC cluster is circled and subsetted for further analysis. **C.** Volcano plot displaying significantly downregulated (skyblue) and upregulated (orange) genes. DE VDCRGs are highlighted and colored (navy blue). **D.** Venn diagram displaying gene counts from this analysis. A total of 11641 genes were detected within the VSMC cluster and tested for differential expression. Of the 11641 genes, 716 genes were downregulated, 2297 were upregulated, and 8628 genes were unchanged. 16/716 downregulated genes and 57/2297 upregulated were VDCRGs. **E.** DotPlot of all DE-VDCRGs in VSMCs displayed as average expression aggregated between conditions (Ctrl vs AAA).

### Vitamin D3 deficient diet exacerbates AAA formation

Using the topical elastase AAA model, wild-type (WT) male mice were maintained on a vitamin D_3_ deficient or sufficient diet for 4 weeks and subsequently treated with either elastase or heat-inactivated (HIE; control) elastase on day 0 (Fig. 2A). Serum analysis of 25(OH)D_3_ (active form of vitamin D_3_) levels confirmed depletion in Vitamin D_3_-deficient mice compared to Vitamin D_3_ sufficient mice (Supplementary Fig. S3). Quantitative analysis revealed a significant increase in mean aortic diameter in the vitamin D_3_ sufficient group compared to controls on day 14 (138.4±8.7% vs. 6.8±1.4%; p<0.0001; Fig. 2B). Importantly, a significant exacerbation in aortic diameter was observed in elastase-treated vitamin D3 deficient mice compared to vitamin D_3_ sufficient group (169±8.5% vs. 138.4±8.7%; p<0.03). Histological analysis of aortic tissue demonstrated that vitamin D_3_ deficient diet led to a significant decrease in smooth muscle alpha-actin (SMα-actin) expression (45.2±4.3 vs. 60.2±4.0%), an increase in elastin fragmentation (221±22.1 vs. 160±10.9 number of breaks/mm^2^), and increased macrophage infiltration (20.2±2.5 vs. 11.2±1.9%) on day 14 compared to mice with vitamin D_3_ sufficient diet (Fig. 2C-F). Furthermore, aortic tissue expression of proinflammatory cytokines (RANTES, MIP-2, MCP-1, CXCL1, and IL-6) and matrix metalloproteinase (MMP2) was significantly increased in WT mice with vitamin D_3_ deficient diet compared to vitamin D_3_ sufficient diet (Fig. 3A-F). A multifold increase in mRNA expression of complement C3 was observed in WT mice with vitamin D_3_ deficient diet compared to vitamin D_3_ sufficient diet (Fig. 3G). There was no difference in mRNA expression of C3aR, C5, and C5aR in the respective cohorts (Supplementary Fig. S4). These findings suggest that vitamin D3 deficiency promotes complement activation at the transcriptional level, potentially contributing to vascular inflammation and AAA progression.

**Figure 2.**
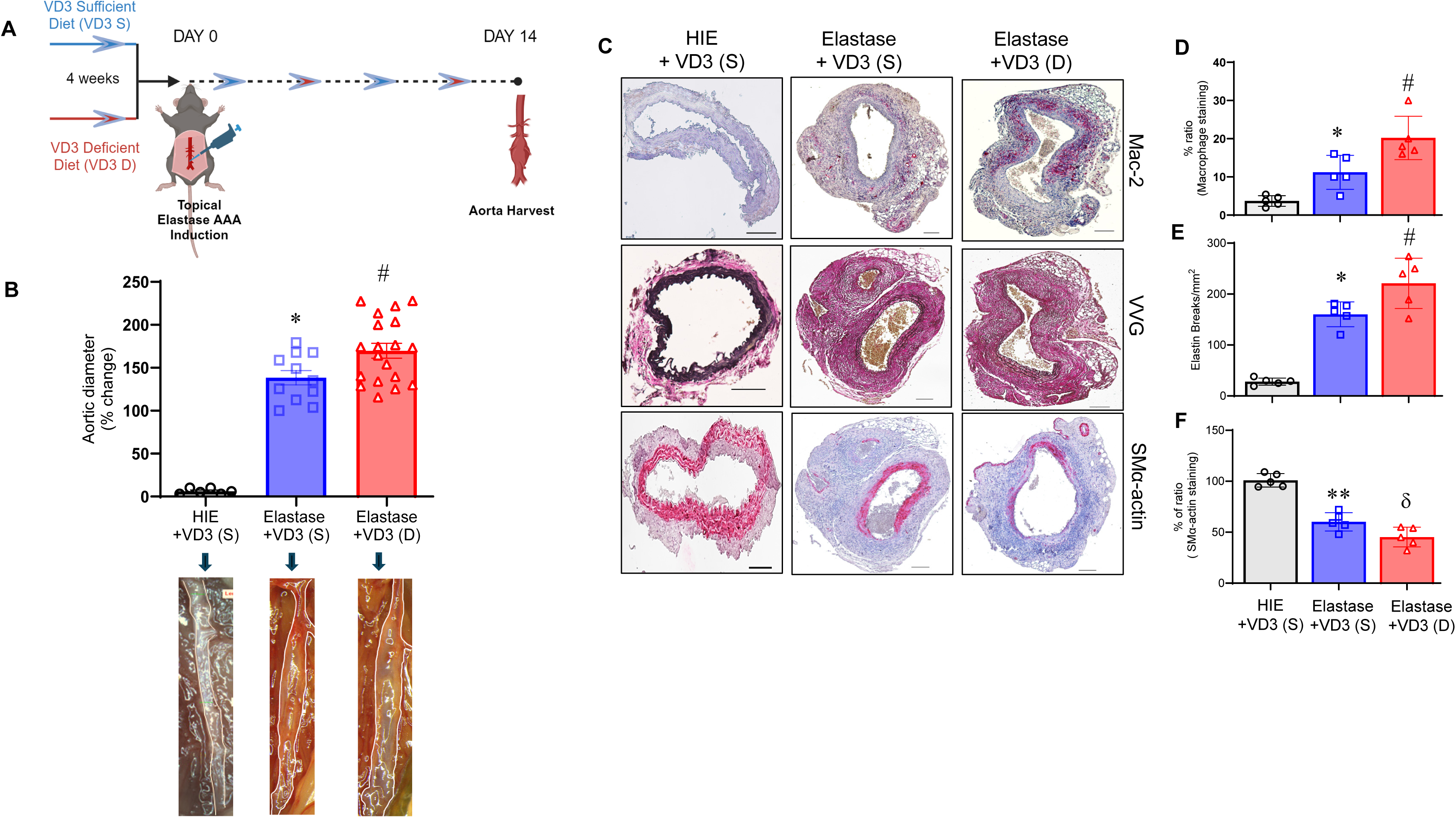
Vitamin D_3_ deficiency promotes AAA formation. **A.** Schematic representation of the experimental design. WT mice were fed either vitamin D_3_–sufficient or –deficient diets starting 4 weeks prior to elastase treatment and aortas were harvested at the endpoint for analysis. **B.** Elastase-treated mice receiving vitamin D_3_–sufficient diet exhibited a significant increase in aortic dilation compared with HIE controls on the same diet. This dilation was significantly exacerbated in mice receiving vitamin D_3_–deficient diet. Representative aortic images from each group are shown. *n* =5–19 mice/group; *p<0.001; ^#^p<0.03. **C–F.** Histological analysis and quantification of aortic sections showed increased macrophage infiltration (Mac-2 staining), disruption of elastic fibers (Verhoeff–Van Gieson staining), and reduced smooth muscle cell α-actin (SM α-actin) expression in VD_3_-deficient compared to VD_3_-sufficient aortic tissue. n=5 mice/group; *p< 0.03; ^#^p<0.01; **p< 0.01; ^δ^p<0.04. HIE, heat-inactivated elastase.

**Figure 3.**
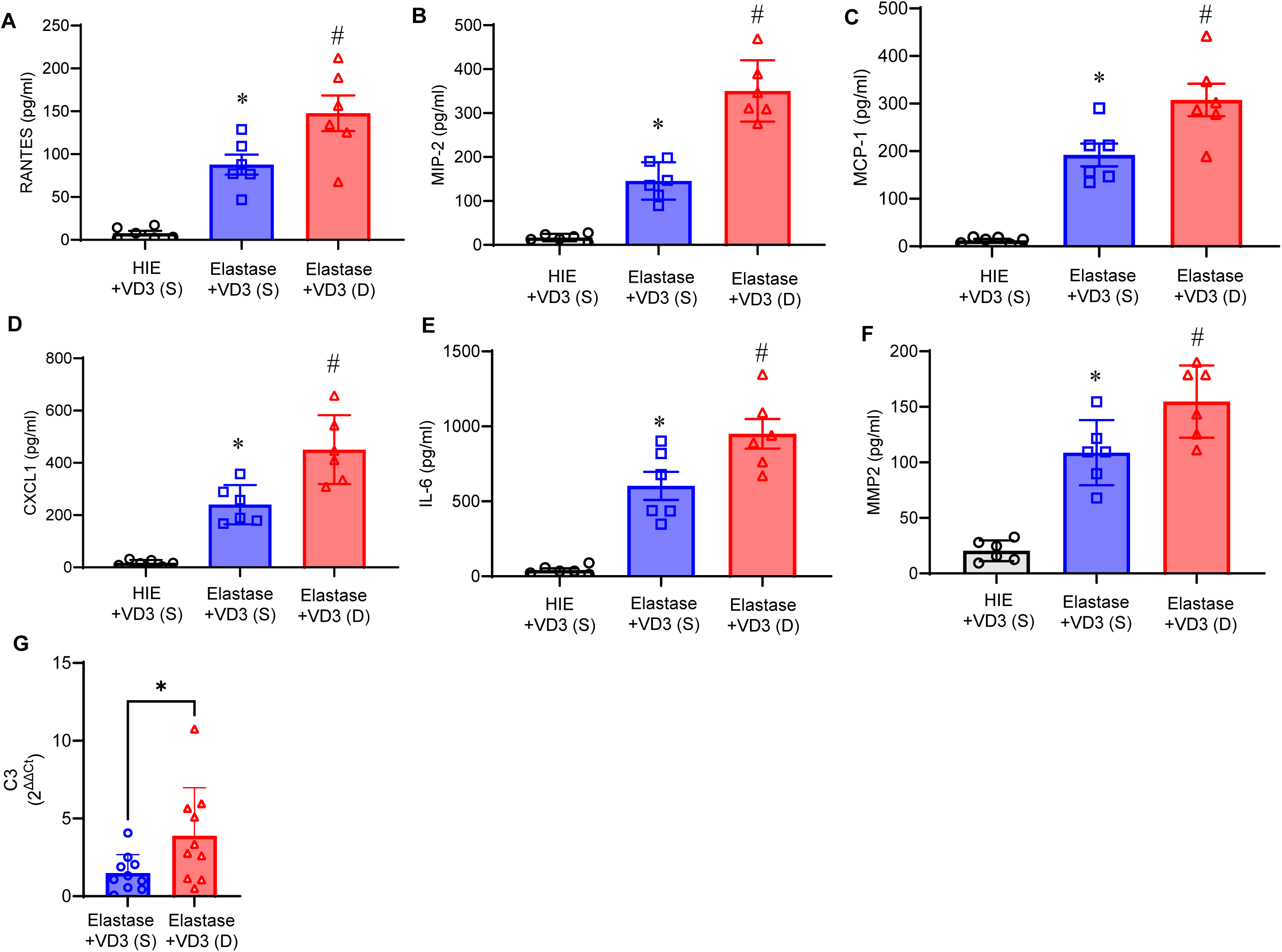
Vitamin D_3_ deficiency enhances aortic inflammation and MMP2 expression. **A-F**. Pro-inflammatory cytokine and MMP2 expressions in aortic tissue of elastase-treated WT mice administered with VD_3_-deficient diet were significantly increased compared to VD3-sufficient mice. *p<0.001; ^#^p<0.04; n=5/group. **G.** mRNA expression of C3 was significantly increased in aortic tissue of elastase-treated VD3-deficient mice compared to VD3-sufficient mice. *p<0.03; n=8-10/group.

### Complement C3aRA treatment attenuates murine AAA formation

Using the topical elastase AAA model, WT male mice were treated with either elastase or heat-inactivated elastase on day 0 and injected with vehicle or C3aRA daily on postoperative days 1 to 13 and harvested on day14 (Fig. 4A). Mice treated with C3aRA demonstrated significantly decreased mean aortic dilation compared to mice treated with vehicle alone (105.2±9.1 vs. 166±15.2%; Fig. 4B). Furthermore, C3aRA treatment led to a significant increase in SMα-actin expression (60.4±3.4 vs. 40.8±5.6%), a reduction in elastin fragmentation (120.0±12.5 vs. 223.8±32.3 number of breaks/mm^2^), and decreased macrophage infiltration (16.4±2.1 vs. 25.4±1.5%) on day 14 compared to vehicle-treated mice (Fig. 4C-F). Moreover, the aortic tissue expression of pro-inflammatory cytokines (RANTES, MIP-2, MCP-1, CXCL1, and IL-6) and matrix metalloproteinase (MMP2) content and activity were significantly attenuated by C3aRA treatment compared to untreated controls (Fig. 5A-F).

**Figure 4.**
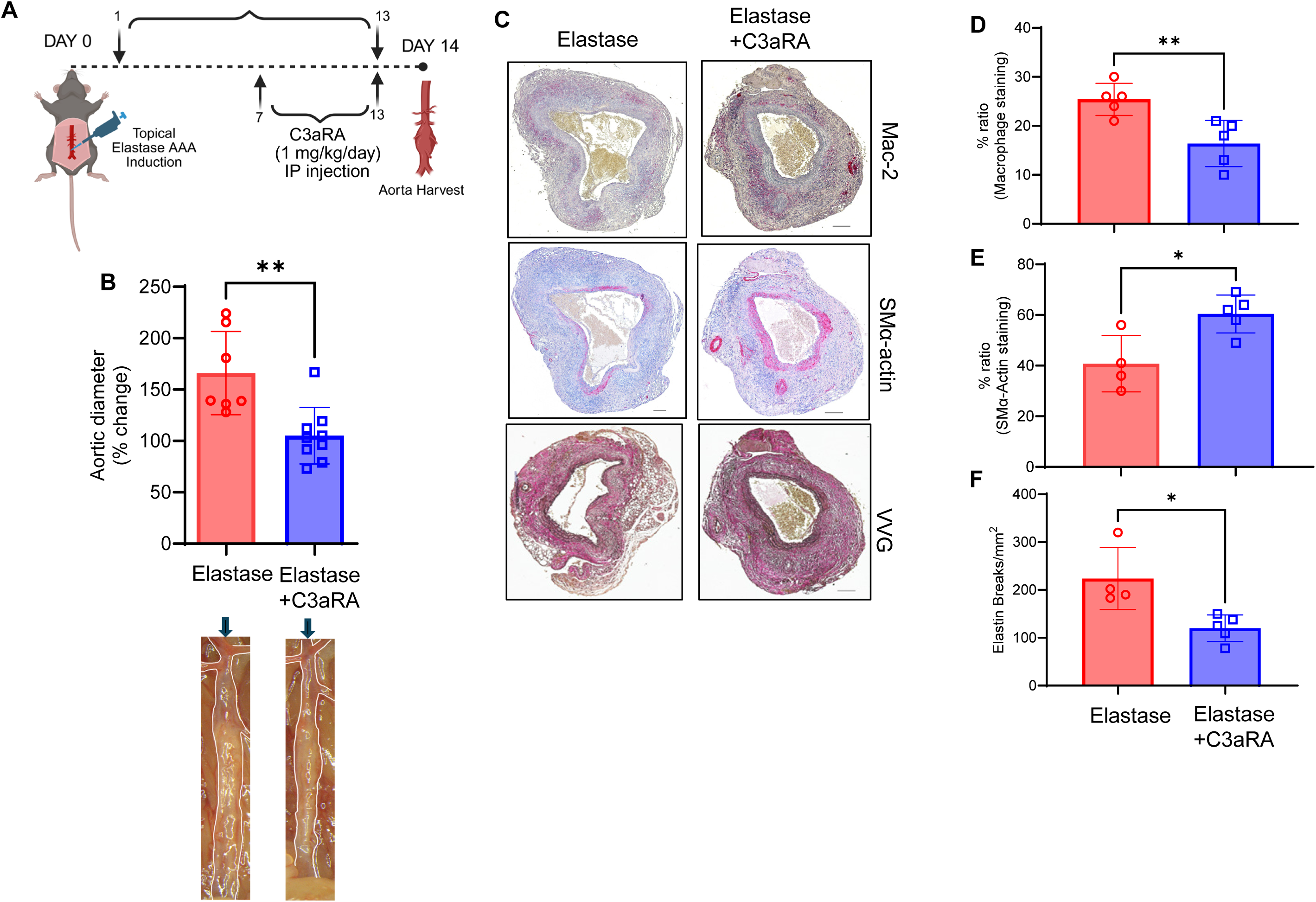
C3aRA treatment attenuates aortic dilation in AAA. **A.** Schematic representation of the experimental design. **B.** Elastase-treated WT mice exhibited significant increase in aortic dilation compared to controls, and that was significantly attenuated by C3aRA administration. Representative images of aortas from each group are shown. **p<0.03; n=7-9/group. **C-F.** Histological analysis and quantification of aortic sections on day 14 demonstrated increased macrophage infiltration (Mac-2 staining), elastic fiber degradation (Verhoeff–Van Gieson staining), and reduced smooth muscle cell α-actin (SM α-actin) expression in elastase-treated aortas. These pathological changes were significantly ameliorated by C3aRA treatment. n=5–10 per group; *p<0.05; **p=0.005.

**Figure 5.**
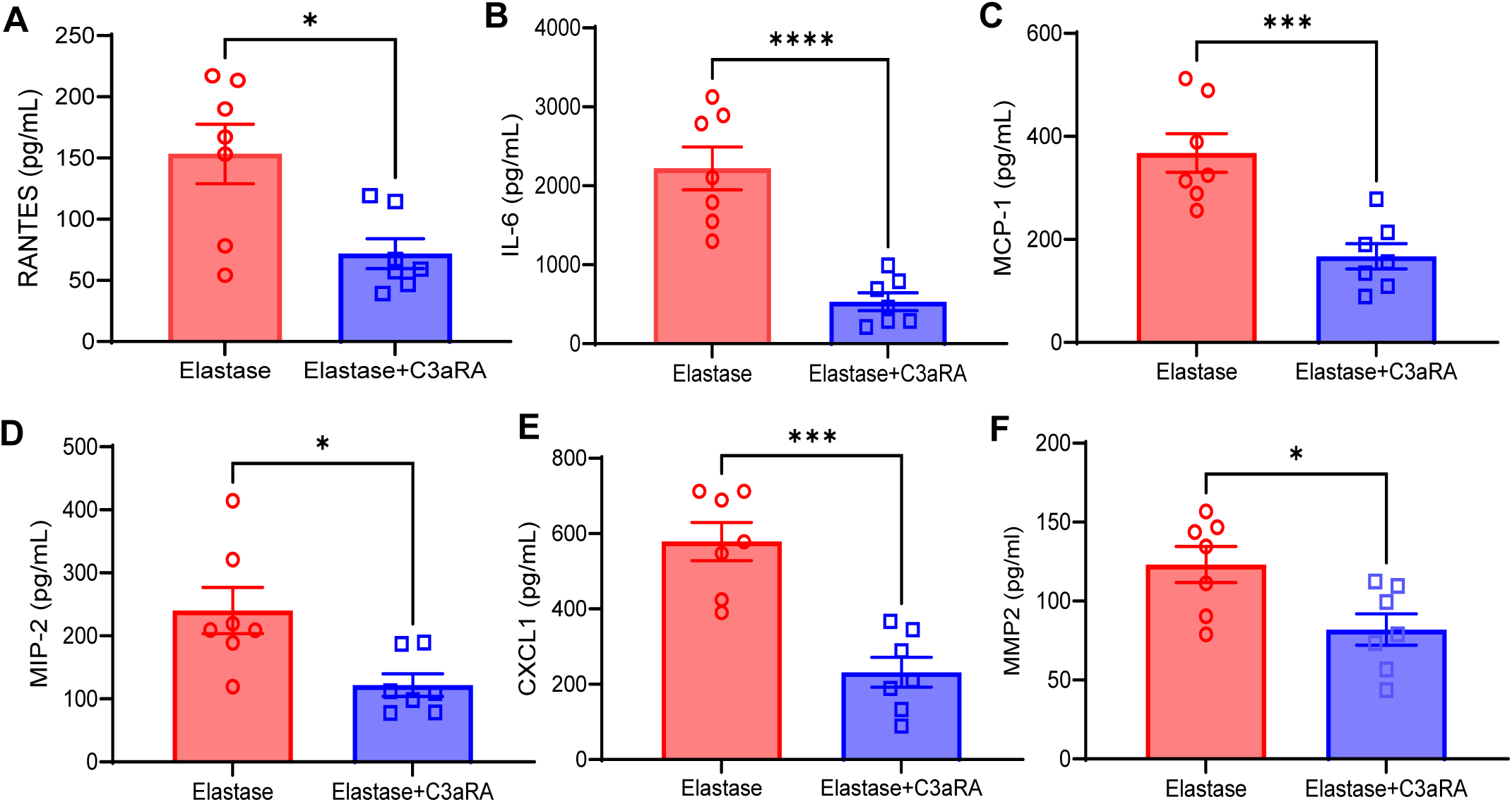
C3aRA administration ameliorates proinflammatory cytokine and MMP2 expression. Proinflammatory cytokine levels **(A-E)** and MMP2 **(F)** expression in aortic tissue were significantly mitigated in the C3aRA-treated group compared to untreated controls. n=7 per group; p < 0.01; *p<0.02;***p=0.001;****p=0.0001.

### Complement C3aRA confers protection against preformed AAAs

Due to the significant reduction in aortic dilation observed with C3aRA administration, we next investigated whether this treatment could also confer protection against already established aneurysms using a chronic AAA and aortic rupture model. C3aRA was administered starting on day 14 post-surgery and continued daily until day 27 and aortic tissue was harvested on day 28 for analysis (Fig. 6A). C3aRA treatment resulted in a significant reduction in aortic dilation compared to saline-treated controls (346.1±13.5 vs. 393±15%, Fig. 6B). Histological analysis and quantitative assessments revealed that C3aRA-treated mice exhibited increased expression of SM-α actin, as well as reduced elastin fragmentation and macrophage infiltration, indicating enhanced vessel integrity (Fig. 6C-F). Furthermore, a significant attenuation of pro-inflammatory cytokine expression and MMP2 content and activity was observed in aortic tissue of C3aRA-treated mice compared to untreated controls (Fig. 6G-J and Supplementary Fig. S5). Collectively, these results suggest that C3aRA treatment mitigates aortic inflammation and vascular remodeling not only during the early phases of AAA development but also in established aneurysms, via mitigation of aortic inflammation and matrix-degrading enzymes leading to preservation of aortic integrity.

**Figure 6.**
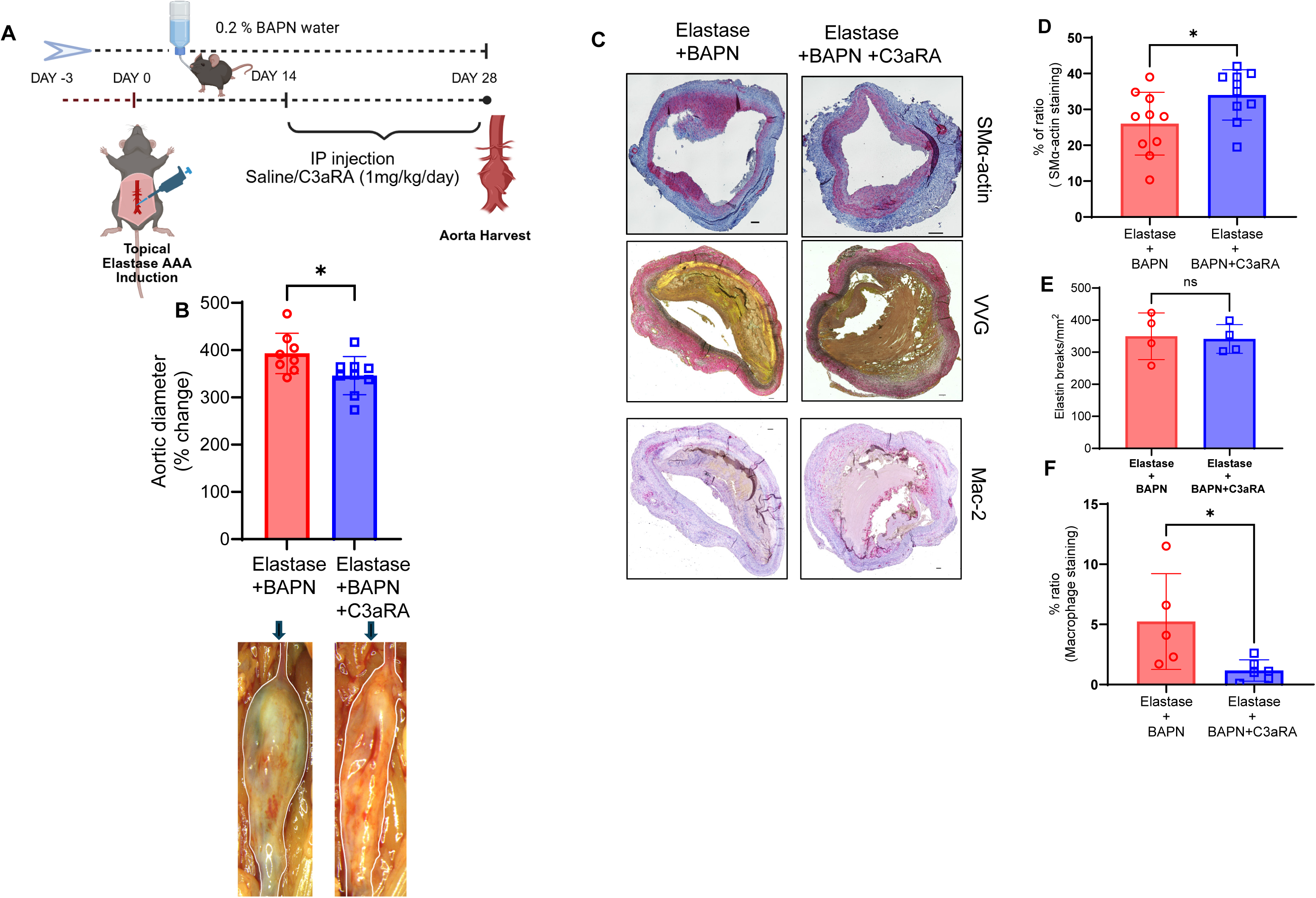
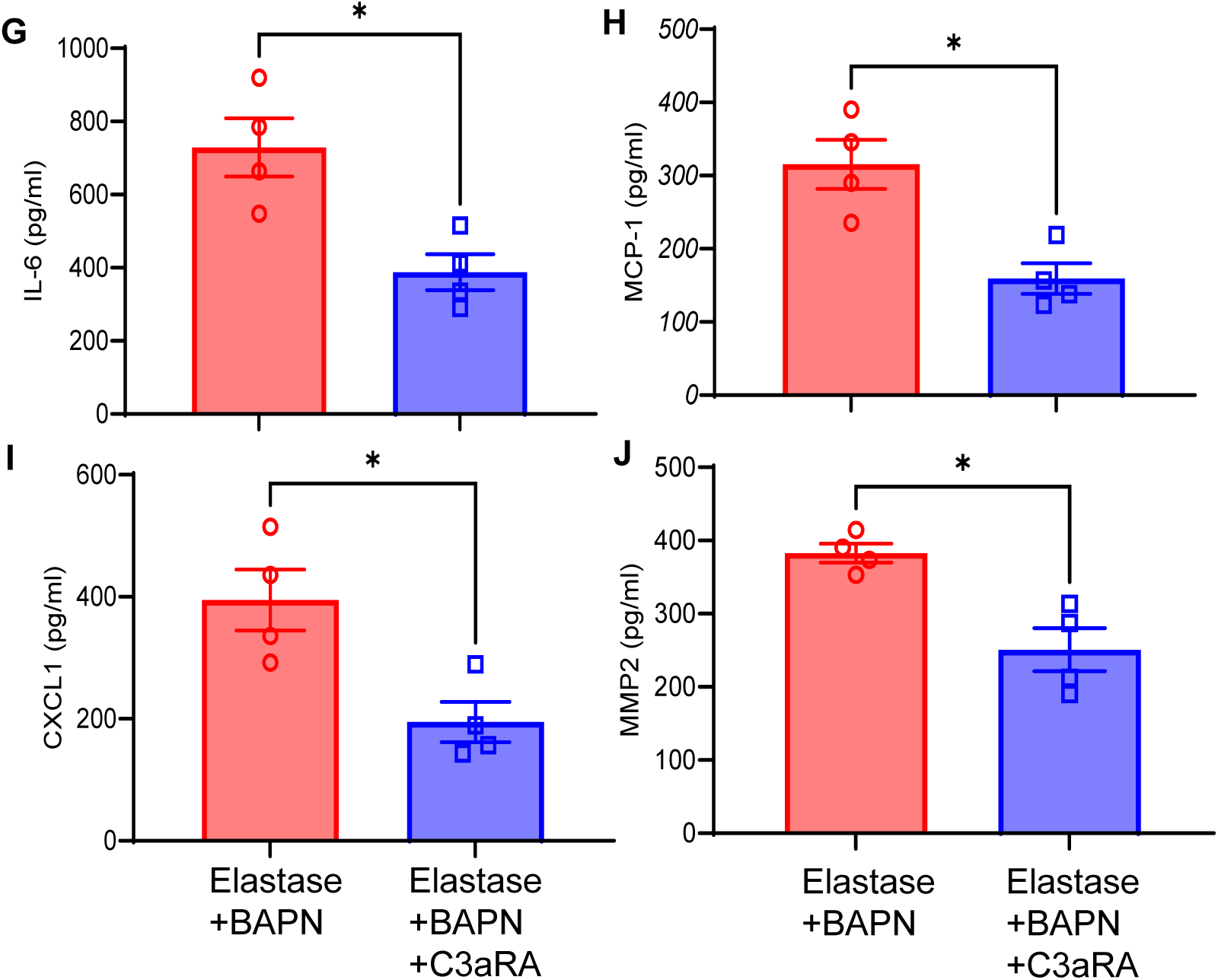
C3aRA administration mitigates preformed AAAs in a chronic murine AAA model. **A.** Schematic of the elastase+BAPN–induced chronic AAA and aortic rupture model. **B.** Aortic diameter on day 28 showed a significant reduction in C3aRA-treated mice compared with elastase + BAPN–treated controls. Representative images of aortic phenotype in each group are shown n = 8–9 mice/group; p<0.05. **C-F.** Representative images and quantification displayed that C3aRA treatment preserved aortic morphology, as evidenced by increased SMα-actin expression, decreased elastin fiber disruption, and reduced macrophage infiltration compared with untreated mice. n=4-10/group; Scale bar=100μm. **G-J**. Treatment with C3aRA significantly mitigated proinflammatory cytokines and MMP2 expression in aortic tissue compared to untreated controls. n=4/group; *p<0.03.

### Vitamin D_3_ Attenuates Elastase-Induced C3a Release in aortic smooth muscle cells

To determine whether treatment with calcitriol (active form of vitamin D_3_) can modulate complement release in vascular smooth muscle cells, we measured C3a levels in HVSMC culture supernatants. Exposure to elastase and cytomix significantly increased the C3a levels when compared to controls (1199±45.8 vs 594±45.2 pg/ml; p<0.001; Fig. 7A-B) indicating robust activation of the complement effector pathway under proteolytic and inflammatory stress. Importantly, treatment with exogenous calcitriol treatment markedly attenuated C3a paracrine release compared to elastase+cytomix group (849.8±25.3 vs 1199±45.8 pg/ml; p<0.001). Furthermore, calcitriol treatment significantly attenuated elastase+cytomix-induced IL-6 secretion and MMP2 expression compared to untreated controls (Fig. 7C-D). These findings demonstrate that calcitriol supplementation can suppresses C3a release by activated HVSMCs, suggesting a direct anti-inflammatory effect on smooth muscle cell–dependent complement signaling during aortic pathologies.

**Figure 7.**
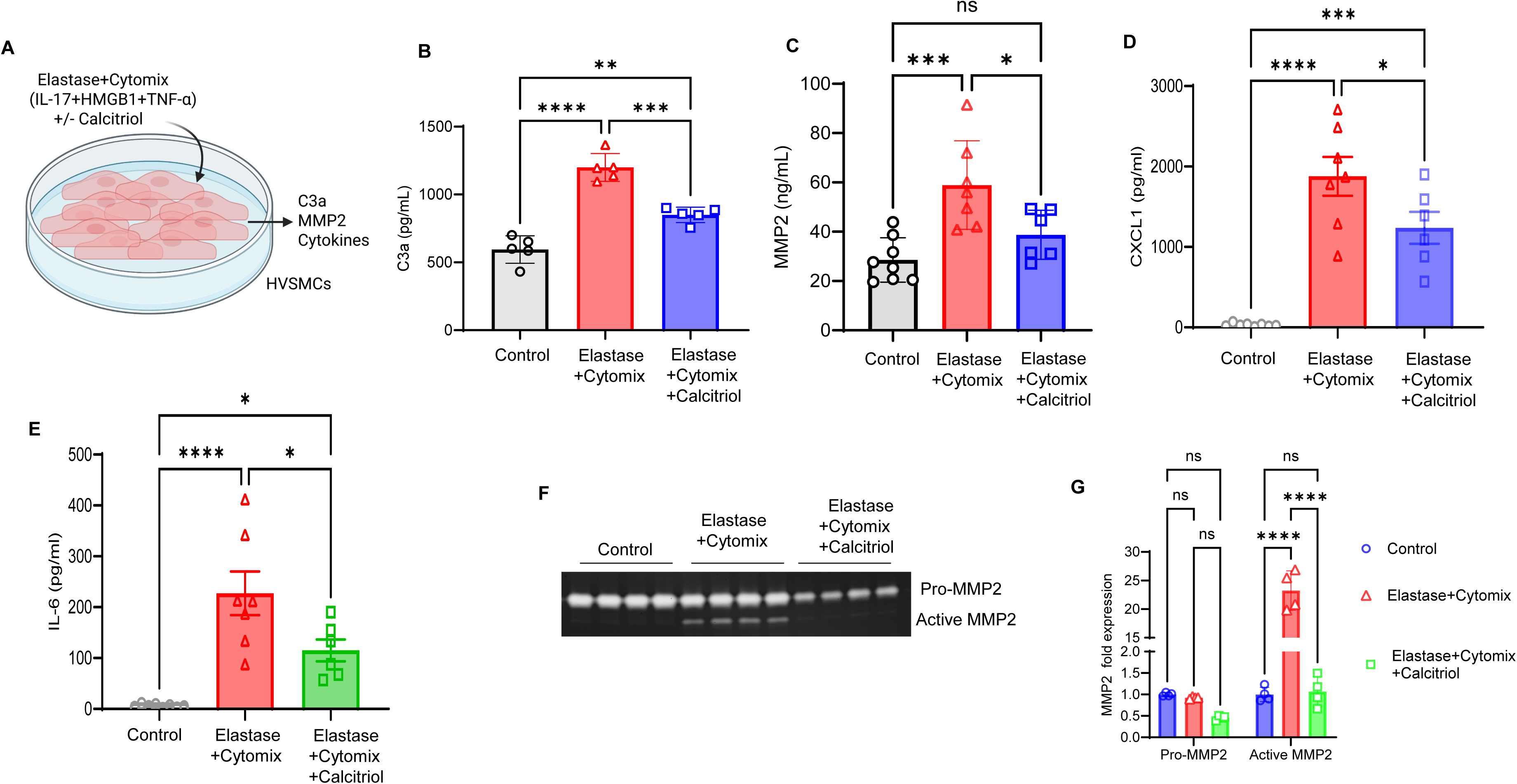
Vitamin D_3_ suppresses complement C3a secretion in human vascular smooth muscle cells. **A.** Schematic for *in vitro* experiment using human primary vascular smooth muscle cells (HVSMCs). Culture supernatants were collected and analyzed for C3a levels by ELISA. **B.** Elastase and cytomix (IL-17A+HMGB1+TNF-α) treatment markedly increased C3a production in HVSMCs compared with controls. VD_3_ administration significantly decreased the expression of C3a in the culture supernatant compared to elastase+cytomix-treated cells. n=5/group. **p < 0.001; ***p=0.0001; ****p < 0.0001. **C-E.** Elastase+cytomix treatment induces a significant increase in MMP2, IL-6 and CXCL1 expression in SMC supernatants, which was attenuated by exogenous calcitriol treatment. n=8/group; *p<0.02; ***p=0.007; ****p<0.0001 ns, not significant. **F-G.** Elastase+cytomix treatment of HVSMCs induces a multifold increase in active MMP2 activity measured by gelatin zymography and quantification, which was significantly attenuated by calcitriol pre-treatment. ns, not significant; ****p<0.0001; n=4/group.

## DISCUSSION

AAA formation is a multifactorial process that involves aortic inflammation and cell death leading to aortic wall weakening, dilation, and eventual rupture. While the exact molecular mechanisms remain elusive, chronic inflammation driven by risk factors such as smoking and hypertension, play a pivotal contributory role in the pathogenesis of progressive aortic dilation and subsequent sudden rupture. While recent studies have implicated an incriminatory role of vitamin D_3_ in AAA development, but the cell-specific molecular mechanistic signaling of remains elusive (12, 13, 24). Our results in human scRNA sequencing analysis demonstrated that vitamin D_3_/complement-dependent genes in SMCs are associated with aortic inflammation, vascular remodeling and cell death. Furthermore, preclinical murine AAA models showed that vitamin D_3_ deficiency increased progression of AAA formation and activation of complement C3 expression, whereas C3aRA treatment mitigates aortic inflammation and vascular remodeling to decrease AAA formation. Finally, *in vitro* stimulation of SMCs showed that vitamin D_3_ supplementation can inhibit C3a release to mitigate IL-6 paracrine secretion and MMP2 content. Taken together, this study provides evidence that vitamin D_3_ deficiency can exacerbate vascular inflammation and remodeling via smooth muscle cell-dependent C3a release to mediate AAA progression and aortic rupture.

Vitamin D_3_ mediated signaling has been shown to immunomodulate the pathobiology of human disease processes, and its deficiency has been linked to chronic inflammatory conditions such as autoimmune disease, cardiovascular disease, and diabetes (25–27). In the context of AAAs, recent findings suggest that relative vitamin D deficiency increases the risk of AAAs, but paradoxically high circulating markers of vitamin D has also been shown to be associated with faster aneurysm growth (28–31). Patients undergoing AAA repair have been reported to be vitamin D insufficient, and a recent study demonstrated that higher circulating 25(OH)D_3_ levels were associated with a lower likelihood of AAA diagnosis (28). Clinical meta-analysis of observational studies have previously reported a significantly lower circulating 25(OH)D_3_ concentration in AAAs compared to controls (11). Similarly, experimental studies using the angiotensin II-infused apolipoprotein E-deficient mouse model, showed that vitamin D supplementation limited AAA growth and reduced rupture rates by modulating proteins involved in extracellular matrix remodeling (12). Similarly, Martorell *et al* showed that calcitriol inhibited endothelial pro-inflammatory and angiogenic chemokine production via vitamin D-retinoid X receptor signaling (13). Importantly, the effects of vitamin D receptor (VDR) activation on endothelial cells can be markedly increased in calcitriol treated mice that can reduce phosphorylation of ERK1/2, p38 MAPK, and NF-κB in ApoE^−/−^ mice. However, the effect of vitamin D_3_ signaling in modulating SMC activation, particularly with complement pathway activation and modulation of MMP2 production and activity, infiltrating inflammatory cells such as macrophages, and degradation of elastin during AAA formation, had to be delineated prior to this study.

In parallel, the activation of complement cascade has been implicated in AAA pathology, as complement activation generates anaphylatoxins C3a and C5a that are known leukocyte chemoattractants and modulators of tissue inflammation (2, 32). The complement system, composed of three main pathways, is a complex primary defense mechanism against pathogenic invasions in human diseases (33). The classical pathway is activated by antibodies bound to antigens in immune complexes, the lectin pathway is activated by mannose-binding lectin, while the alternative pathway can be activated by the spontaneous hydrolysis of C3(32). All these pathways can lead to inflammation, promote antibody production, assist in phagocytosis, and attack cell membranes. Human tissue analyses have demonstrated increased C3a, C3b, C3AR1, and C5AR1 expression in AAA samples compared with controls (32). Complement C3/C3a has been shown to be increased in plasma and aortic tissue of aortic dissection patients with a potent macrophage-SMC axis driving the pathology, wherein C3a deposition and activation of SMCs is observed (34). Additional studies have shown increased deposition of C3, IgG, and chemotactic mediators, such as RANTES, in aneurysm tissue (35, 36). While few studies have examined the interplay between vitamin D3 and complement activation in AAA formation, the role of complement in aneurysm development has been previously reported (2, 32, 36, 37). Complement involvement has also been demonstrated in elastase-induced murine AAA, where alternative pathway activation promotes aneurysm formation via C3a and C5a, and mRNA levels of the membrane receptors C3AR1 and C5AR1 were also significantly increased in AAA tissue (2). These receptors are important mediators of the complement cascade components that can induce inflammation, a characteristic feature of the AAA tissue. Our study provides converging *in vivo* and *in vitro* evidence that vitamin D modulates AAA formation through regulation of complement activation, particularly the C3a axis. Also, treatment with C3aRa, but not C5aRa mitigated experimental AAA formation (data not shown). Additionally, vitamin D_3_–deficient diet significantly increased C3 expression, enhanced aortic dilation, as well as hallmarks of aortic inflammation (proinflammatory cytokine expression) and vascular remodeling (MMP2 expression), leading to loss of smooth muscle cell integrity, and increase in elastin degradation and macrophage infiltration.

Although the contribution of vitamin D metabolism in vascular pathologies have been shown to activate immune cells and activated T cells within the aortic wall during the degenerative process of aortic aneurysms, here our studies add complement component C3a as an additional regulator of this excessive inflammation and remodeling (5, 38). However, there are several limitations to the current study. While our results suggest that vitamin D_3_ deficiency exacerbates the progression of AAAs and aortic ruptures by enhancing C3 activation, extracellular matrix degradation and enhanced inflammation, the multifactorial pathology may also be affected by concurrent alterations in calcium, fibroblast growth factor 23 (FGF23), phosphorus, and parathyroid hormone (PTH) relative to serum concentrations of 25(OH)D_3_. Serum concentrations of calcium, PTH, FGF23 and phosphorus, that are known biomarkers in the vitamin D metabolic pathway, can potentially contribute to AAA development as they have been linked to several risk factors, such as hypertension, vascular inflammation and calcification (39). This association with vitamin D metabolism is further complicated by sex specific differences in AAA pathology as higher calcium levels in women, but not men, can double the risk of AAA formation independent of cardiovascular risk factors. Conversely, we did not investigate if C3aRA treatment had any impact on VDR expression. Further exploration of the key metabolites of vitamin D pathway are required to decipher the chronicity of AAA pathology relative to calcium signaling in a sex-dependent manner.

In summary, our study identifies a previously underappreciated regulatory pathway linking vitamin D_3_ homeostasis to complement activation in AAA. Vitamin D_3_ deficiency promotes complement expression and immune cell infiltration to compromise aortic wall integrity and enhance aneurysm growth. Pharmacological C3aR antagonism further validates this pathway as a therapeutic target to mitigate progression of AAAs and prevent aortic rupture. Together, these findings establish the vitamin D_3_-complement axis as a key regulator of AAA pathogenesis and a potential avenue for future intervention.

## Acknowledgments

The authors would like to thank Tabitha Randi for her impeccable help with administrative duties and laboratory management.

## Supplementary Material

Supplementary material (Figures S1-S5 and Table 1) are available online.

## Author contributions

AA and JH contributed equally to the manuscript. A.A., J.H., W.U., J.A.V., G.S., M.F., S.S., C.A., J.M., and A.K.S. conducted the research, performed experiments, collected, analyzed and/or interpreted data. A.A., A.K.S., and G.R.U. drafted and edited the manuscript. A.K.S. and G.R.U. conceived and designed the study, supervised the project, obtained research funding and have access to all the data. All authors read the final manuscript and gave final approval of the version submitted.

## Data Availability

All relevant data are included in the manuscript or the Supplementary material.

