## Supplemental Figures 1-5 for "Vitamin D3 Deficiency Exacerbates Abdominal Aortic Aneurysm Progression Via Complement C3a Activation"

### Slide 1
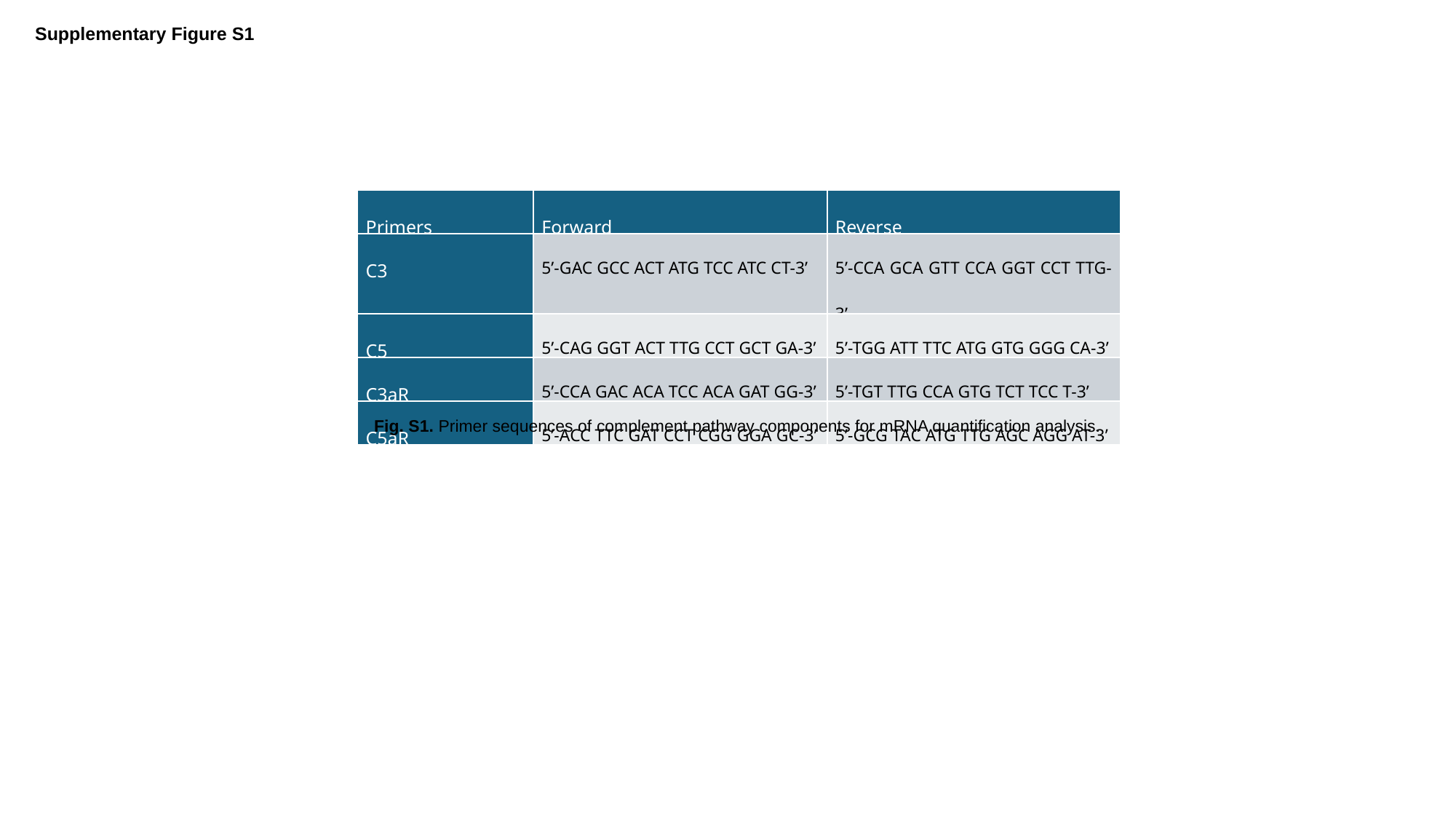

Supplementary Figure S1
| Primers | Forward | Reverse |
| --- | --- | --- |
| C3 | 5’-GAC GCC ACT ATG TCC ATC CT-3’ | 5’-CCA GCA GTT CCA GGT CCT TTG-3’ |
| C5 | 5’-CAG GGT ACT TTG CCT GCT GA-3’ | 5’-TGG ATT TTC ATG GTG GGG CA-3’ |
| C3aR | 5’-CCA GAC ACA TCC ACA GAT GG-3’ | 5’-TGT TTG CCA GTG TCT TCC T-3’ |
| C5aR | 5’-ACC TTC GAT CCT CGG GGA GC-3’ | 5’-GCG TAC ATG TTG AGC AGG AT-3’ |
Fig. S1. Primer sequences of complement pathway components for mRNA quantification analysis.

### Slide 2
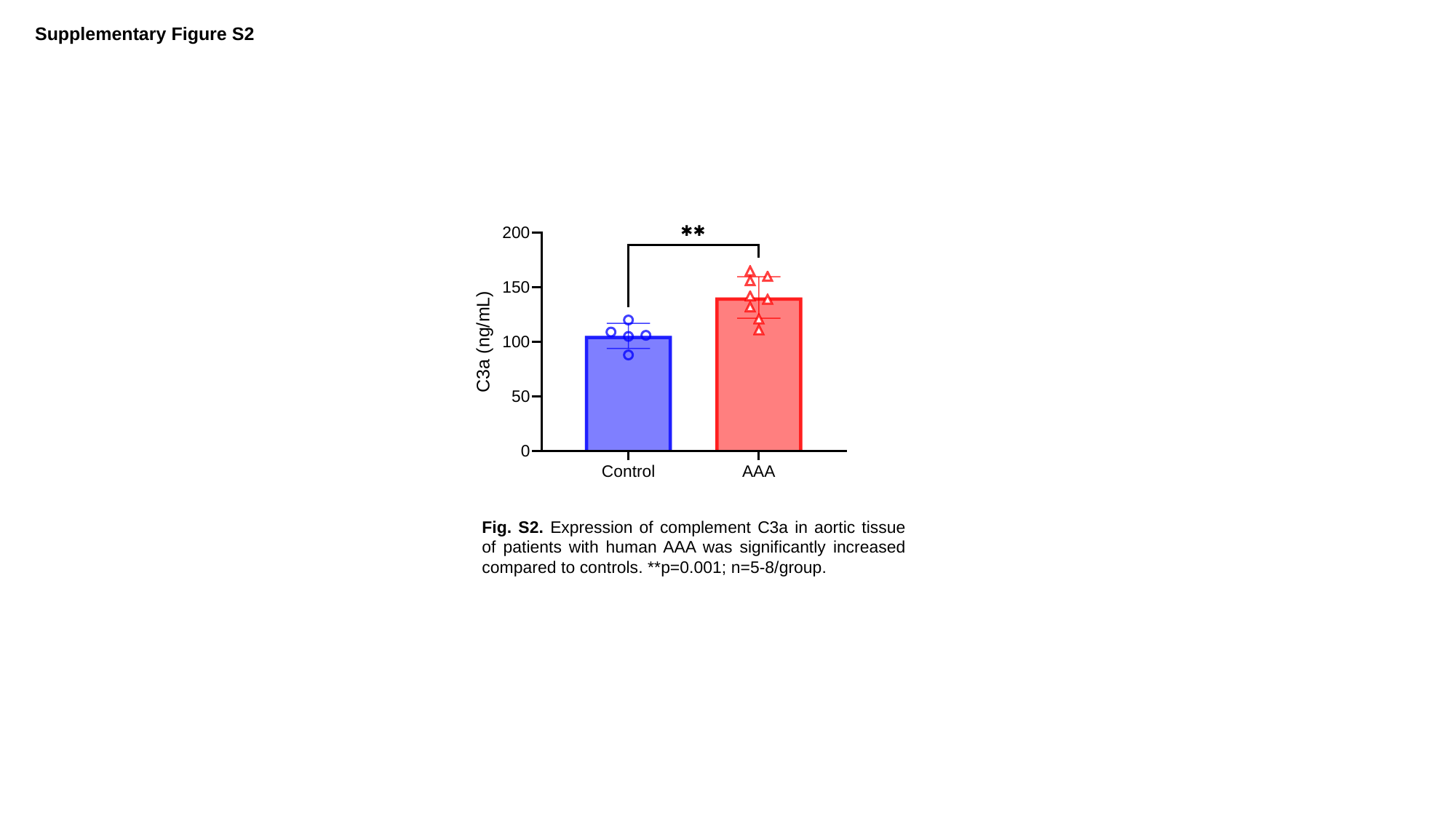

Supplementary Figure S2
Fig. S2. Expression of complement C3a in aortic tissue of patients with human AAA was significantly increased compared to controls. **p=0.001; n=5-8/group.

### Slide 3
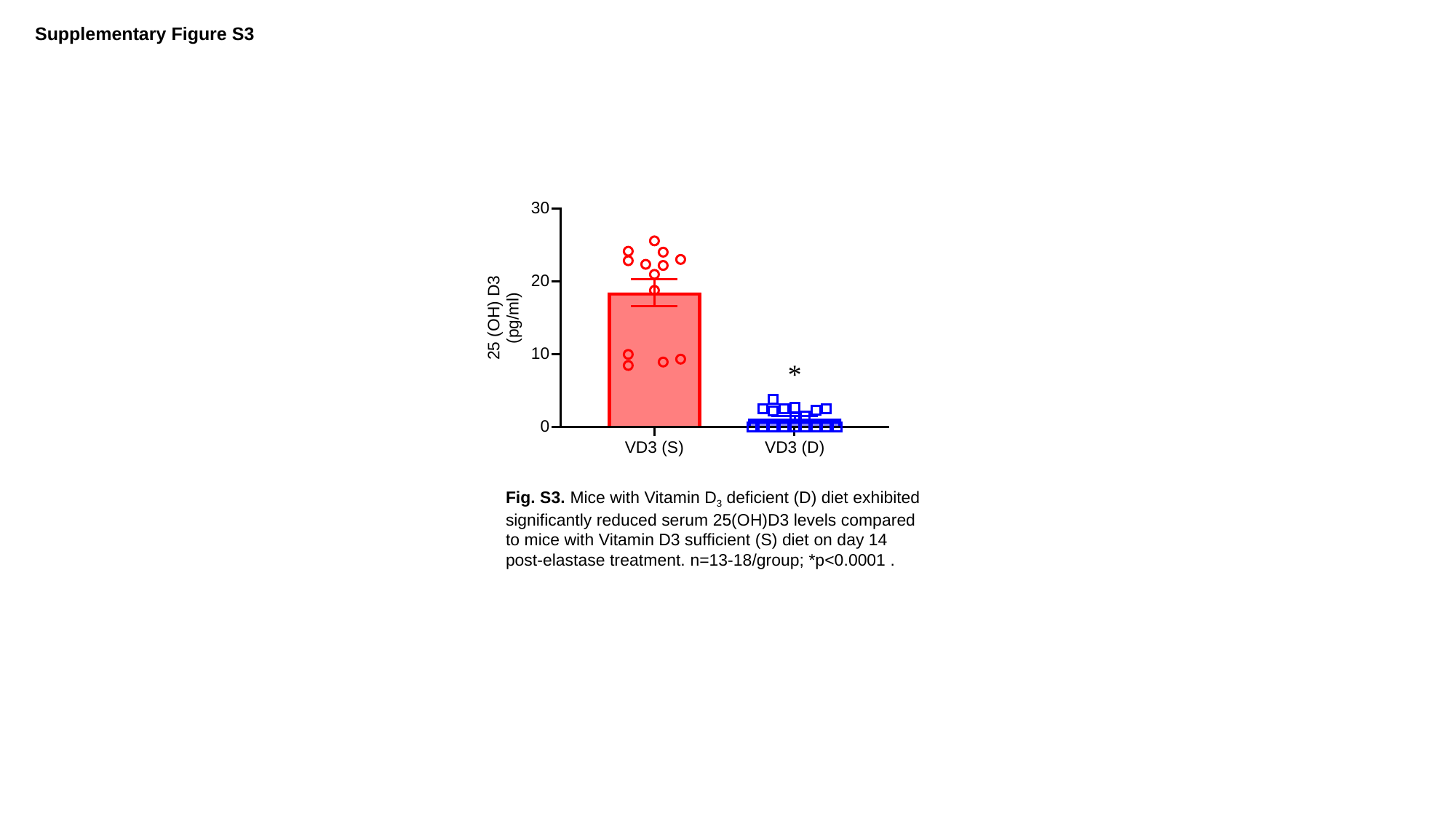

Supplementary Figure S3
*
Fig. S3. Mice with Vitamin D_3_ deficient (D) diet exhibited significantly reduced serum 25(OH)D3 levels compared to mice with Vitamin D3 sufficient (S) diet on day 14 post-elastase treatment. n=13-18/group; *p<0.0001 .

### Slide 4
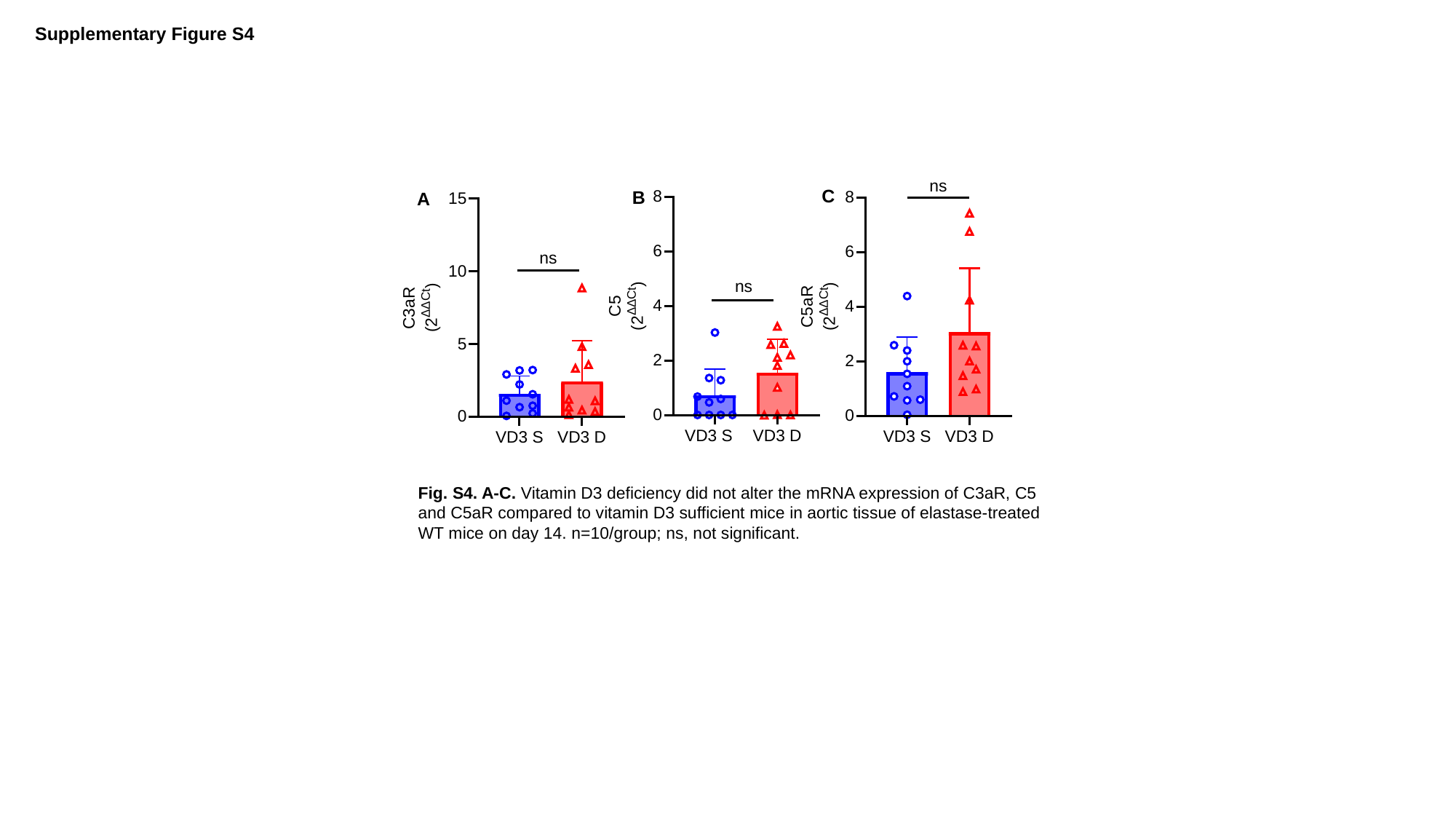

Supplementary Figure S4
ns
C
B
A
ns
ns
Fig. S4. A-C. Vitamin D3 deficiency did not alter the mRNA expression of C3aR, C5 and C5aR compared to vitamin D3 sufficient mice in aortic tissue of elastase-treated WT mice on day 14. n=10/group; ns, not significant.

### Slide 5
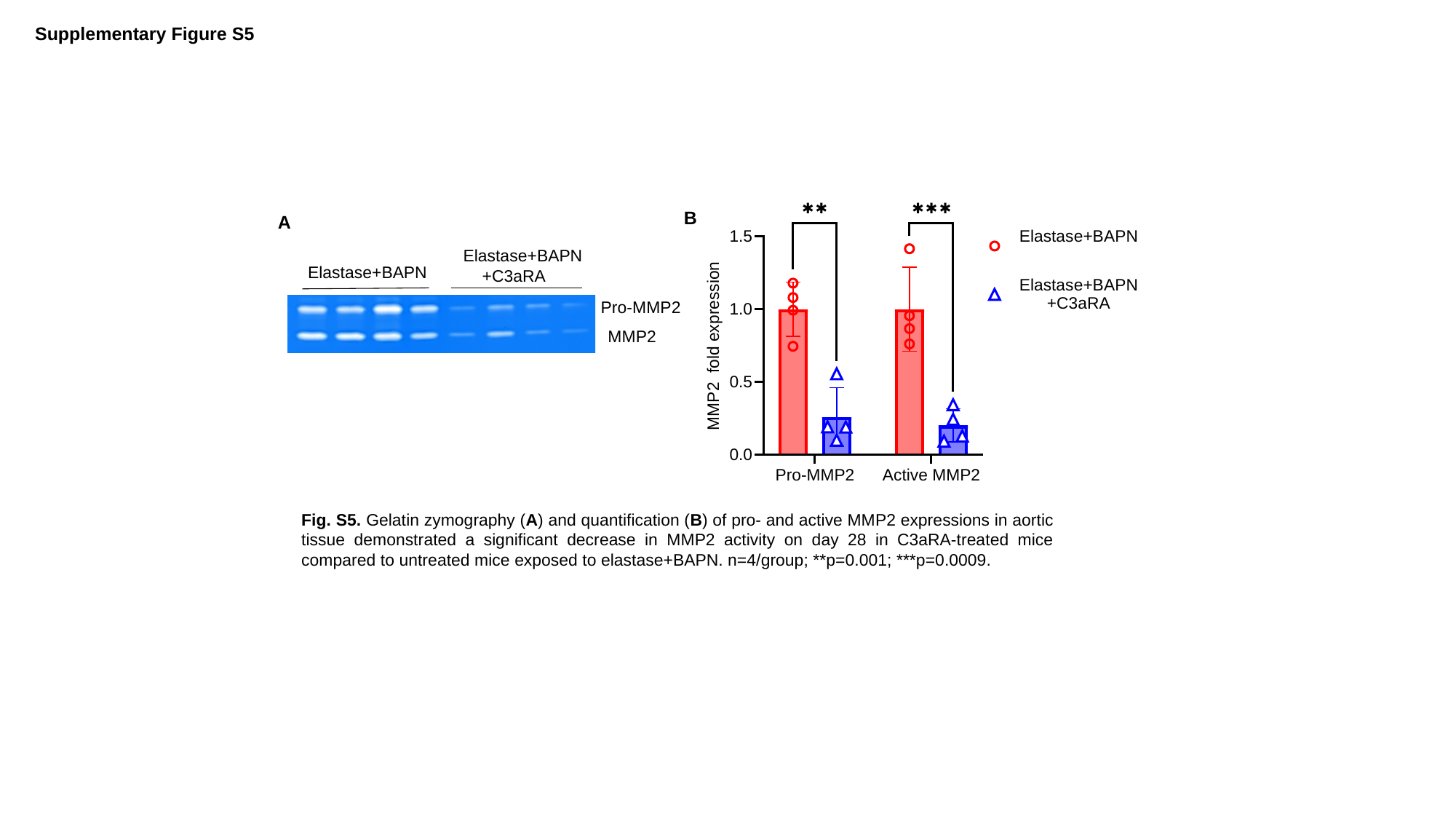

Supplementary Figure S5
B
A
Elastase+BAPN
 +C3aRA
Elastase+BAPN
Pro-MMP2
 MMP2
Fig. S5. Gelatin zymography (A) and quantification (B) of pro- and active MMP2 expressions in aortic tissue demonstrated a significant decrease in MMP2 activity on day 28 in C3aRA-treated mice compared to untreated mice exposed to elastase+BAPN. n=4/group; **p=0.001; ***p=0.0009.
